# Development and optimization of the host DNA depletion in blood cultures using a saponin and SAN nucleases-based method

**DOI:** 10.1101/2025.07.14.664698

**Authors:** Jawad Ali, Anurag Basavaraj Bellankimath, Jonathan Hira, Crystal Chapagain, Rafi Ahmad

## Abstract

Bloodstream infections and sepsis are major health issues causing millions of deaths annually. Early and accurate diagnosis of sepsis is crucial for reducing mortality and combating antimicrobial resistance. However, current diagnostic methods are slow and time-consuming. Nanopore sequencing has the potential to serve as a rapid diagnostic method for sepsis. However, extracting bacterial DNA from blood samples is challenging due to the high content of host DNA. This study aimed to develop and optimize a method using saponin and SAN nucleases for effective host DNA depletion in blood cultures. Different concentrations of saponin, HL-SAN, and M-SAN nucleases, combined with various salt conditions (NaCl and MgCl2), were tested for their ability to deplete host DNA in blood cultures spiked with *E. coli* and *S. aureus*. The impact of different bead beating durations on DNA fragment sizes was also examined. The efficiency of host DNA depletion and bacterial DNA extraction was assessed using quantitative PCR (qPCR) and nanopore sequencing. Results indicated that 4% of saponin effectively lysed host cells, releasing DNA while preserving bacterial cells. The host DNA is then efficiently depleted using 250 units (10 μL) of HL-SAN and M-SAN nucleases. Although SAN in combination with higher salt concentrations (2.5 M NaCl and 50 mM MgCl2) showed somewhat better host DNA removal, the difference was not statistically significant. Reducing bead beating time to 6 minutes improved the recovery of longer DNA fragments compared to 10 minutes. These results demonstrate an effective method for host DNA depletion using saponin and SAN nucleases, which is compatible with downstream processes, such as nanopore sequencing and qPCR.

## Introduction

Bloodstream infections (BSIs) and sepsis are major health problems accounting for millions of deaths each year around the world [1]. Early and appropriate diagnosis of sepsis is critical for the patient’s survival. However, the current sepsis diagnosis is based on traditional blood culture followed by biochemical methods, MALDI-ToF, PCR, and usually automated antibiotic susceptibility testing (AST) [2]. Although these methods are considered the gold standard for sepsis diagnosis, they are time-consuming and may take up to 72 hours for pathogen identification and antibiotic resistance [3]. The delay is undesirable for septic patients, as they require prompt administration of antibiotics. Therefore, there is an urgent need to develop rapid diagnostic methods for sepsis.

Multiplex PCR systems overcome some limitations of traditional blood culture methods because they are quick and can deliver results within 1 to 4 hours after a positive blood culture. However, multiplex PCR panels are limited to identifying only a certain species and resistance markers and need regular updates to include new resistance patterns [4–7]. Metagenomics next-generation sequencing (mNGS) using nanopore technology (Oxford Nanopore, UK) is currently one of the most promising technologies for rapid diagnosis of infections. mNGS overcomes the limitations of multiplex PCR as it can detect any pathogen or resistance markers in the sample and does not rely on predefined targets. Nanopore sequencing provides real-time data, low cost per sample, and the potential to be used directly on samples without the need for culturing [8,9]. Several studies have demonstrated the potential of nanopore sequencing for detecting BSI-causing pathogens and their antibiotic resistance genes (ARGs) [10–13]. Nanopore sequencing of the positive blood cultures has been shown to identify bacterial species with 97% sensitivity and 94% specificity in 3.5 hours, compared to the routine diagnostic methods [10]. Also, the AMR results for the top ten sepsis-causing pathogens were obtained 20 hours earlier compared to the routine antibiotic susceptibility testing (AST) [10]. Similarly, *Harris et al*. have also demonstrated that nanopore sequencing of the positive blood cultures provided a concordance of 94% for species identification compared to the routine diagnostic methods [11]. A recent study has used nanopore-targeted sequencing, utilizing pathogen-specific primers for multiplex PCR, and identified 114 pathogens associated with BSIs in a few hours [12]. *Moragues-Solanas et al*. have shown that nanopore sequencing of the spiked blood samples with sepsis-relevant pathogens can identify the bacteria and antibiotic resistance determinants in less than 12 hours [14]. In our previously published study, we have shown that nanopore sequencing can identify sepsis-causing pathogens and their antibiotic resistance genes (ARGs) in seven to nine hours [15].

Although nanopore sequencing shows promise for rapid BSI diagnosis, it requires the extraction of high-molecular-weight bacterial DNA. However, DNA sequencing from blood samples faces several challenges. The number of circulating microbes in BSIs is usually low (1 – 100 CFU/mL), and the host DNA from white blood cells significantly exceeds the microbial DNA [16,17]. Additionally, the human genome comprises approximately 6 billion bases, whereas the average bacterial genome is roughly 3.5 million bases [18,19]. In a one-to-one comparison of the genome content, the total nucleic acid amount in a bacterium is only about ∼0.07% of the DNA in a single mammalian cell. Due to these complexities, extracting sufficient quantities of bacterial DNA from blood samples for detection by nanopore sequencing remains a significant challenge.

To address the challenge of high human and low microbial DNA content in clinical samples, several studies have employed the selective lysis of host cells using nonionic detergents, such as saponin, Triton X-100, and Tween-20 [20–24]. Specifically, saponin, which has a specificity for cholesterol-rich plasma membranes, has been explored for the selective lysis of host cells in clinical samples, such as blood, urine, and sputum [23,25–28]. This process of selective host lysis releases host DNA, which is then removed in a subsequent step. Chemical methods, such as propidium monoazide, and enzymatic methods based on endonuclease enzymes (like DNase I/II, benzonase, and salt-activated nucleases), have been used to deplete host DNA before total DNA extraction for downstream applications [21,29,30]. Recently, heat-labile salt-activated endonucleases (HL-SAN) and medium salt-activated endonucleases (M-SAN) have been studied for digesting the released host DNA in sputum and UTI samples [26,28,31]. HL-SAN is optimized for digesting nucleic acids in high salt conditions. The added salt also enhances chromatin disruption, which accelerates host depletion by this enzyme. [32,33]. Meanwhile, its counterpart, M-SAN, is designed to deplete nucleic acids under physiological salinity conditions, facilitating the detection of a wide range of bacterial and viral pathogens. [33]. The concentration of saponin, enzymes, and the optimal conditions for enzyme activity are essential for the selective and effective depletion of host DNA. While the concentration of saponin has been studied for effective selective lysis of host cells in blood, limited research has focused on optimizing SAN concentrations and salt conditions needed for maximum endonuclease activity [23].

In this study, we aim to optimize host DNA depletion in blood cultures by testing different concentrations of saponin and SAN enzymes, as well as various salt conditions (NaCl and MgCl2) to enhance enzyme activity. Previous studies have evaluated HL-SAN exclusively for host DNA digestion in blood cultures [14]. However, the present study is the first to assess the performance of M-SAN for removing host DNA from blood culture samples. Additionally, we evaluated different bead beating (BB) times for bacterial DNA extraction to recover longer DNA fragments suitable for nanopore sequencing. We demonstrated that 4% saponin is optimal for the selective lysis of host cells, followed by digestion of the released DNA using 10 μL (250 U) of SAN enzymes. The different salt concentrations tested did not significantly affect enzyme activity. However, it is recommended to use 2.5 M NaCl and 50 mM MgCl2 as buffers for these enzymes in clinical samples. A shorter BB time of 6 minutes was more effective in recovering longer DNA fragments suitable for nanopore sequencing compared to a 10-minute BB.

## Methods

### Bacterial strains

In this study, two of the most common bacterial pathogens that cause sepsis, *Escherichia coli* and *Staphylococcus aureus,* were used for testing. The *E. coli* isolate, identified as NCTC 13441, was obtained from the National Collection of Type Cultures (NCTC), Public Health England. Similarly, *S. aureus* CCUG 17619 was obtained from the Culture Collection of the University of Gothenburg (CCUG, Sweden). *E. coli* NCTC 13441 is an Extended-spectrum beta-lactamase (ESβL) positive resistant isolate that has been tested in our previously published studies [15,34]. However, the *S. aureus* CCUG 17619 isolate is wild type with no phenotypic resistance reported. These bacterial strains were stored at -80 °C in glycerol and were refreshed one day before the experiments on agar plates.

### Testing of saponin concentration for selective lysis of host cells and preservation of bacterial cells

A range of saponin concentrations was tested to achieve blood cell lysis without affecting the viability of the bacterial cells. Bacterial growth dynamics were employed to test the preservation of bacterial cell viability following saponin treatment. Thus, bacterial growth dynamics were measured using the standard micro dilution method. Before measuring growth kinetic parameters, bacterial strains were prepared according to the following steps.

Bacterial strains were revived from glycerol stock and isolated as single colonies from an overnight static growth culture on Brain Heart Infusion (BHI) agar medium (VWR Life Sciences). Individual colony from each strain was inoculated in 3 mL of BHI broth and cultivated for 2-3 hours with moderate shaking at 37 °C, depending on the strain’s background. After brief cultivation, the culture samples were centrifuged at 5000 rpm for 10 minutes. The supernatant was discarded, and the pellets were resuspended in Phosphate-Buffered Saline (PBS) (VWR Life Sciences). Bacterial cell density was adjusted to 10^5^ CFU/ml and seeded into each well of a 96-well plate at the final concentration. Saponin was added to the wells at a final concentration of 2-5%, with increments of 1%. The Breathe-Easy sealing membrane (Z380059, Merck) was used to seal the plates, which were then incubated at 37°C in a Synergy H1 microplate reader. Absorbance at 600 nm for each well was measured at an interval of 30 min over 24 hours (Figure 1A).

**Figure 1.**
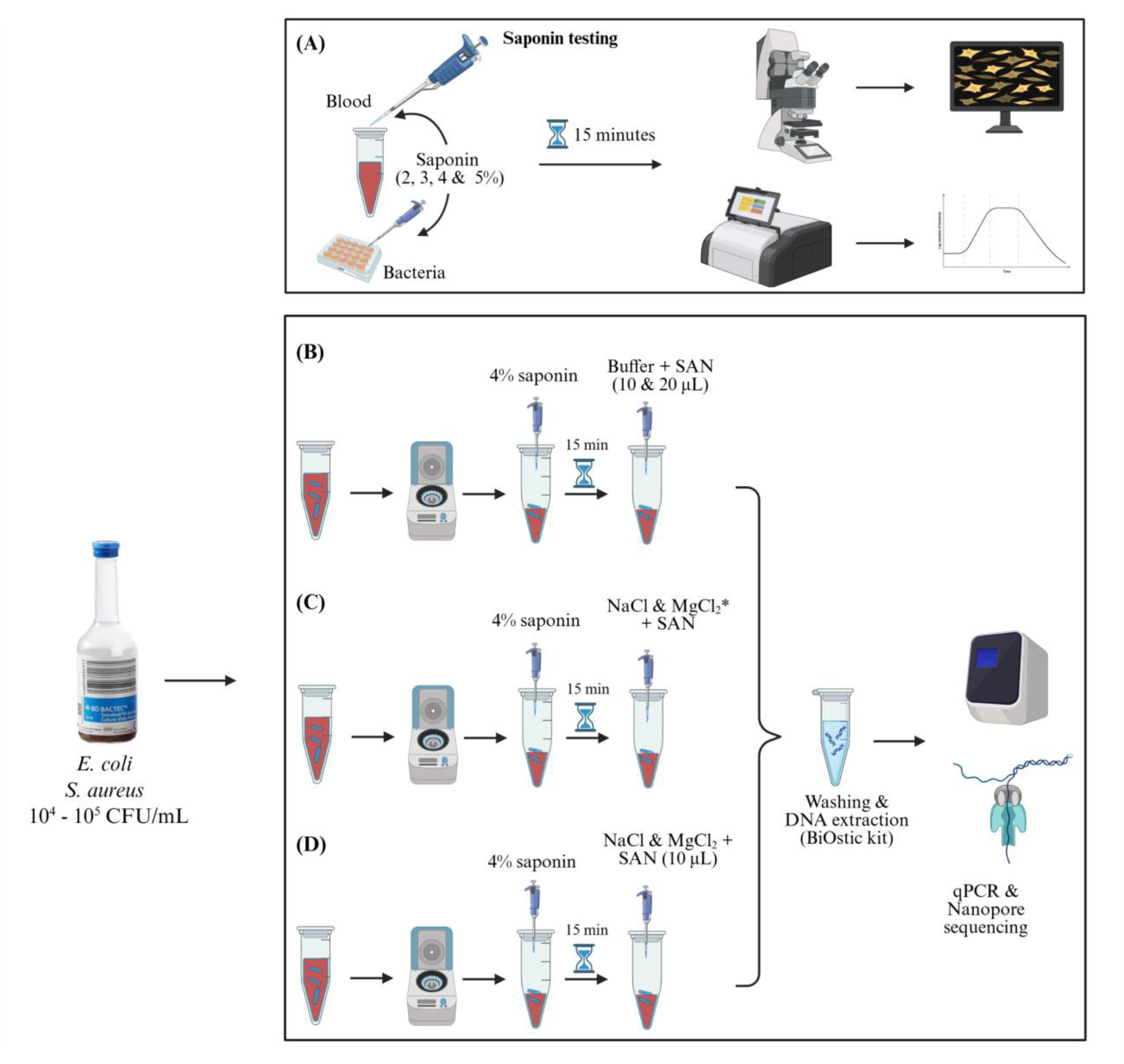
An overview of the experimental design. (A) Testing different concentrations of saponin (2, 3, 4, and 5%) to see the lysis of host cells and bacterial cells. (B) Host DNA depletion of *Escherichia coli* and *Staphylococcus aureus* blood cultures using saponin and different concentrations of SAN, followed by DNA extraction. (C) Testing the effect of different salt concentrations on the activity of SAN. (D) The final optimized protocol using 4% saponin, 10 μL (250 U) of SAN, 2.5 M NaCl, and 50 mM MgCl2 for effective host DNA depletion. * 2 vs 2.5 M NaCl + 50 mM MgCl2, 2.5 M NaCl + 15 vs 50 mM MgCl2, 0.5 M NaCl + 15 mM MgCl2 vs 2.5 M NaCl + 50 mM MgCl2.

### Growth data analysis using QurvE

Acquired growth data was analyzed using QurvE software v1.1. Raw data generated from the microplate reader was transformed to fit in QurvE, and parameters such as growth OD threshold, time at 0 hour (t0), and maximum time for growth (tmax) were set while quality check on raw growth curve. In this study, the growth threshold was set to 1.5, and tmax to 24 hours to extract growth kinetics parameters. Growth profiling of individual bacteria was performed using a growth-fitting exponential growth model with a heuristic linear regression on log-transformed data. Growth rates (µ) were measured for individual bacteria cultivated in growth medium with saponin and without saponin (PBS as negative control). A linear fit was performed using an R2 threshold of 0.95, a Relative standard deviation (RSD) of 0.1, and a dY threshold of 0.05 to estimate the difference in maximum growth and minimum growth. The growth rate measured with the linear fit model was validated with the Gompertz parametric fit model. Additionally, the growth kinetics parameter doubling time was measured on log-transformed data using a spline fit model, with the smoothing factor set to 0.55. The static growth of bacteria was qualitatively validated using the droplet growth method. After 10-15 minutes of incubation at room temperature, 10 µl droplets with 3 replicates from each saponin concentration were plated on BHI-agar plates and incubated overnight at 37 °C. Initial stock concentrations were ∼10^5^ CFU/mL. Any countable colonies >30 from a 10 µl droplet will confirm the non-detrimental effect of saponin on bacterial growth. On the other hand, brightfield imaging was performed briefly to monitor the lysis of blood cells as supporting evidence of host cell depletion. Brightfield image of a blood sample treated with a targeted saponin concentration acquired using a microscope (Nikon Eclipse Ti) at 200x magnification.

### Preparation of bacterial inoculum and blood culturing

The bacterial isolates were revived from the glycerol stocks by streaking on BHI agar medium. The bacterial inoculum was prepared as previously described [15] with minor modifications. Briefly, three to four colonies from the overnight-grown bacterial cultures were suspended in fresh BHI broth medium and incubated for two hours. After incubation, the cultures were centrifuged, and the supernatant was discarded. The bacterial pellet was then resuspended in PBS, and the absorbance was measured at an optical density (OD) of 600 nm using a spectrophotometer (Thermo Electron Corporation, BioMate 3, USA). Dilutions of these suspensions were performed to achieve 1000 CFU/mL, and 1 mL was added to 10 mL of defibrinated sheep blood (ThermoFisher Scientific). Along with spiking the sheep blood, these suspensions were cultured on agar plates and incubated overnight to determine the colony-forming units per mL (CFU/mL). The spiked blood samples were then inoculated into BD BACTEC™ Plus Aerobic medium (BD) culture bottles and incubated at 37 °C. Bacterial growth in the blood culture was monitored by drawing blood samples after each hour of incubation and plating them on BHI agar plates. The goal was to obtain blood cultures with 10^4^ – 10^5^ CFU/mL. In addition to culturing the blood samples on BHI agar plates, aliquots of the samples were also stored at -20 °C. Those that were confirmed to have 10^4^ – 10^5^ CFU/mL were later used for DNA extraction.

### Optimization of SAN nucleases for host DNA depletion

#### Concentration of the SAN nucleases

The host DNA depletion in blood cultures was performed by treating the blood cultures with 4% saponin, followed by the addition of HL-SAN and M-SAN (ArcticZymes Technologies, Norway). Before the addition of endonucleases, a specific buffer consisting of 2.5 M NaCl and 50 mM MgCl2, with pH levels of 7 and 8.3, was added for M-SAN and HL-SAN, respectively. In our previous study, we used 20 μL (500 U) of enzyme to deplete the host DNA in urine samples obtained from patients with urinary tract infections (UTIs) [24]. In this study, we used two different concentrations, 10 μL (250 U) and 20 μL (500 U) of both HL-SAN and M-SAN, to assess their performance for host DNA depletion in blood cultures. The idea was to determine if lower concentrations of SAN nucleases could achieve comparable host depletion efficiency to higher concentrations (Figure 1B).

#### Testing various salt concentrations for SAN activity

After optimizing the endonuclease concentrations, various salt combinations were tested to create the buffer for enzymes, assessing their impact on enzyme activity and host DNA depletion. Initially, the buffer was made in nuclease-free water at a concentration of 2.5 M NaCl and 50 mM MgCl2, as previously reported [24]. The enzyme activity was evaluated for host depletion using this buffer and then compared with a lower concentration of 2 M NaCl, while maintaining a consistent concentration of MgCl2 (50 mM).

Next, the concentration of NaCl in the buffer was maintained at 2.5 M, and two different concentrations of MgCl2 (15 mM and 50 mM) were tested. Lastly, a different buffer combination, using 0.5 M NaCl and 15 mM MgCl2, was also tested and compared with the other previously tested salt concentrations. All these different combinations of salts were tested for their ability to deplete host DNA by SAN in both *E. coli* and *S. aureus* blood cultures (Figure 1C). The efficacy of the enzymes for host DNA depletion at various salt concentrations was assessed using either qPCR, sequencing, or both, as described in the following sections.

### DNA extraction and quality assessment

After the selective depletion of host DNA using saponin and SAN endonucleases, DNA was extracted from all the samples. The QIAamp BiOstic Bacteremia Kit (Qiagen, Germany) was used for DNA extraction, as previously described [15,35]. Briefly, the DNA pellet obtained after selective host depletion was treated with the lysis buffer from the BiOstic kit, and the manufacturer’s protocol was followed to extract DNA. Along with the DNA extractions from host-depleted samples, a subset of samples was also extracted using the standard BiOstic kit protocol, without the host depletion step with saponin and SAN.

The quantification of all the extracted DNA samples was performed using the Qubit 4 fluorometer with Qubit 1X dsDNA HS Assay Kit (ThermoFisher Scientific). The purity of the extracted DNA was evaluated using a Nanodrop ND-1000 spectrophotometer (ThermoFisher Scientific), which measured the absorption ratios at wavelengths of 260/280 nm and 260/230 nm. DNA fragment size distribution and integrity were evaluated using the Agilent 4150 TapeStation System with the Genomic DNA ScreenTape assay (Agilent Technologies, USA).

### Optimization of the bead beating protocol

The BiOstic kit utilizes a combination of chemical and mechanical methods to lyse cells and release DNA. The manufacturer’s default protocol suggests ten minutes of continuous bead beating after adding lysis buffer to the sample. In these experiments, we used three cycles of 2 minutes of bead beating, with a 1-minute incubation on ice between each cycle. The BB was performed using a VWR Vortex laboratory shaker (444-2790) at 2500 rpm (15,708 rad.min-1). The samples extracted using the reduced bead beating protocol were compared with those from the default protocol to assess any changes in DNA fragment lengths (as determined by TapeStation) and their impact on the sequencing output.

### qPCR analysis

All the extracted DNA samples were evaluated for the relative proportions of host (sheep) and bacterial DNA. For the quantification of bacterial and host DNA, species-specific primers were used (Supplementary Table 1). The qPCR reactions were performed using a 7500 Fast Real-Time PCR system (Invitrogen™, USA). Each PCR reaction consisted of 3 μL 5X HOT FIREPol^®^ EvaGreen^®^ qPCR Supermix (Solid BioDyne, Estonia), 0.2 μM of forward and reverse primers, 10.4 μL of nuclease-free water, and 1 μL of template DNA. The conditions for qPCR amplification were as follows: initial denaturation at 95 °C for 12 min, 40 cycles of 95 °C for 25 seconds, 60 °C for 45 seconds, and 72 °C for 60 seconds, followed by a dissociation stage. The assessment of host DNA depletion and bacterial DNA enrichment was performed using the ΔCt values, which were normalized to the Ct of the reference control samples. The normalized Ct values were converted to fold changes using the 2^-ΔCt^ method. Statistical analysis was performed using a 2-tailed unpaired t-test to assess the significance between the different experimental groups.

### Nanopore sequencing and bioinformatics analysis

Some of the extracted DNA samples were sequenced utilizing nanopore sequencing (Oxford Nanopore Technologies, UK). The sequencing was performed using the MinION MK1D device with an R10.4.1 flow cell (FLO-MIN114). Library preparation for sequencing was performed using the Rapid Barcoding Kit 96 V14 (SQK-RBK114.96) and the Rapid PCR Barcoding Kit 24 V14 (SQK-RPB114.24). The Rapid PCR Barcoding Kit was used for samples with low DNA concentrations that were not compatible with the input requirements of the Rapid Barcoding Kit.

Raw sequencing data were collected using the ONT MinKNOW GUI software (Version 6.0.11), and fast basecalling was performed in real time through the Dorado basecaller. The unclassified reads were recovered using an in-house build tool, MysteryMaster (Abdolrahman Khezri, Sverre Branders, JA, ABB, CC, Manfred Grabherr, RA). The identification of bacterial pathogens from the sequencing data was performed as described previously [24]. Briefly, the basecalled sequencing data were BLAST searched against a custom reference database of the most common sepsis-causing pathogens. Reads that did not align with any prokaryotic genome were presumed to be the host reads. To evaluate the presence of mitochondrial DNA percentage in the host DNA, raw reads were also BLAST searched against the Ovis aries (sheep) mitochondrial genome from the NCBI database.

## Results

### 4% saponin showed selective lysis of host cells while preserving bacterial cells

The results indicate that the control exhibited the highest planktonic growth rate as measured by growth rate μ with a linear fit model and 95% CI. Gradually, the growth rate decreased as saponin concentrations increased, which demonstrates a dose-dependent inhibition (Figure 2). At 2-4% saponin, *E. coli* and *S. aureus* exhibit similar growth trajectories, which suggests minimal differences across these concentrations. However, a noticeable reduction in planktonic growth was observed in 5% saponin, indicating the initiation of an inhibitory effect at this higher concentration. Growth trends observed in the growth curve also indicate that the growth rate increases over time, peaking between 4 and 6 hours for *E. coli*. The control sample represents the topmost peaks, followed by saponin-treated samples, which reinforces the saponin effect on bacterial growth. On the other hand, *S. aureus* displays a sharp decrease in growth rate at the beginning of saponin treatment, but steadily increases from 4 hours, indicating cellular adaptability and viability in the context of dysbiosis events. However, the control samples again show the highest peak after 6 hours and then stabilize after 16 hours for *S. aureus*. Doubling time measured for both *E. coli* and *S. aureus* supports the growth rate trajectories exerted by saponin. The lowest doubling time was recorded for samples with no saponin treatment, whereas the highest doubling time was observed for 5% saponin-treated samples. Contrary to this, even though no saponin demonstrated the lowest doubling time, samples with 5% saponin had the second lowest doubling time, and this might be due to viability dysbiosis. Additionally, static growth events in the presence of saponin also demonstrate minimal inhibitory activity on static cellular viability. Qualitative droplet-based static growth strategies indicate steady growth of bacterial colonies on BHI agar, except for a difference observed for *S. aureus* at 5% saponin, where a smaller number of colonies were recorded. This suggests that saponin levels above 5% may be detrimental to bacterial growth. On the other hand, microscopic examination of saponin-treated blood samples also supports the selectivity of saponin towards host cells rather than bacteria at given concentrations. In this study, 4% saponin was considered for evaluation, and therefore, blood samples that undergo saponin treatment demonstrate lysis of red blood cells (RBCs) and white blood cells (WBCs), whereas platelets remain intact (Supplementary Figure S1). The presence of mitochondrial host DNA was also evaluated, and the results showed that the sequencing data only contained 0.1% mitochondrial DNA. The saponin and SAN-based method, along with the chromosomal DNA, also effectively depleted the host mitochondrial DNA. Overall, *E. coli and S. aureus* seem to tolerate saponin concentrations of 2-4% quite well, while allowing blood cells, except platelets, to be completely lysed. To keep the downstream experimental workflow active, 4% saponin is considered a targeted saponin based on its selective host blood cell lysis and its support for both static and planktonic growth of bacteria.

**Figure 2.**
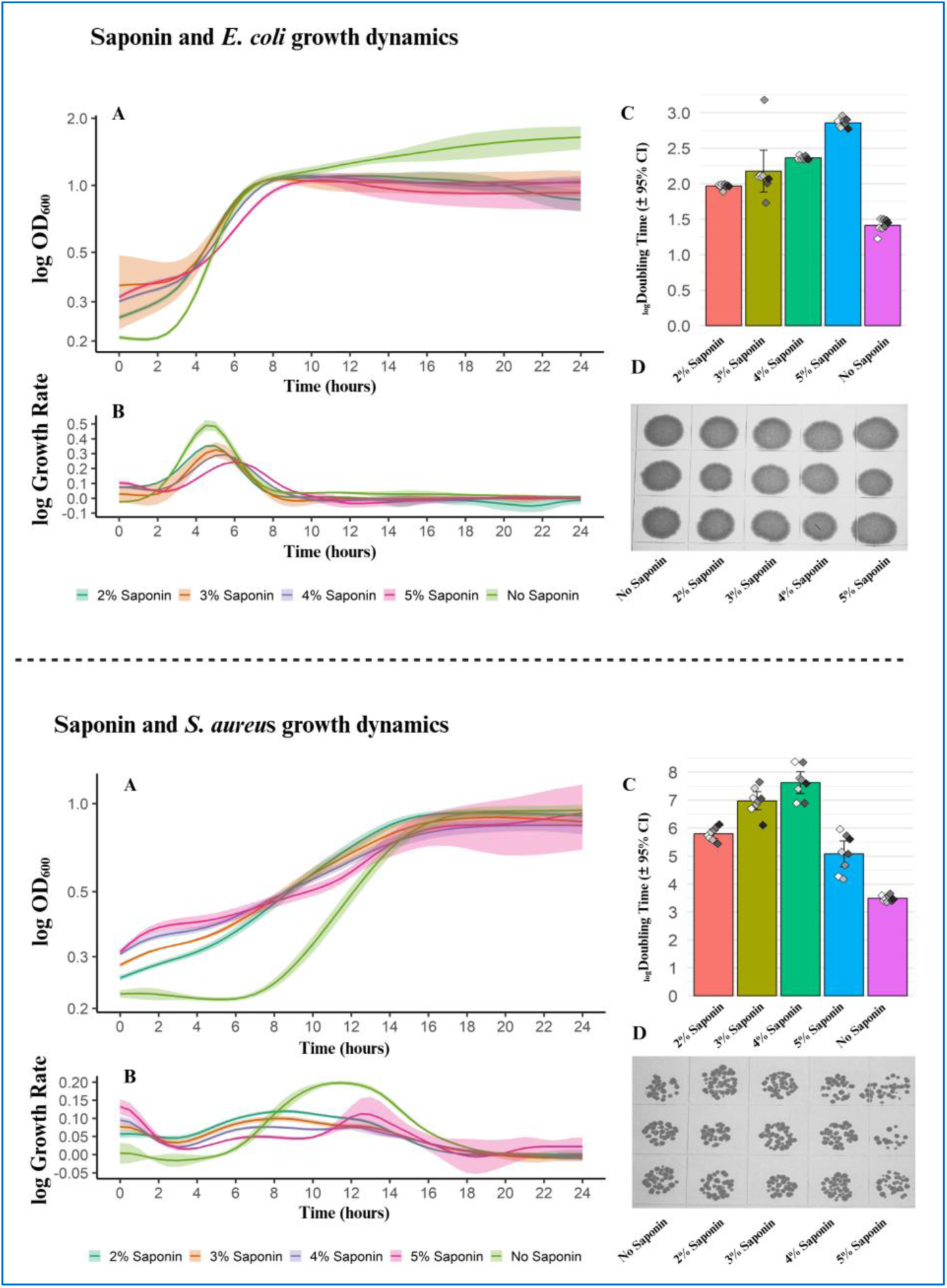
**Effect of saponin on bacterial growth dynamics (*Escherichia coli* and *Staphylococcus aureus*).** For both strains (A) Growth curves over time (24 hours) for conditions with Control (PBS), 2-5% saponin. Growth curves depicting log-transformed OD600 on the y-axis and hours on the x-axis. (IB) Smoothed line histogram demonstrating the distribution of growth rates across saponin conditions, with overlapping curves indicating variability. (C) Doubling time (log transformed) highlighting differences in growth kinetics. Bar plot with (CI-95%) for each condition depicting gradual changes in doubling across the saponin concentration. (D) Static growth images on BHI agar for both strains illustrating no effect of saponin on colony formation on agar.

### Blood cultures achieved 10^4^ – 10^5^ CFU/mL after 2 – 4 hours of incubation

To optimize the bacterial inoculum for blood culturing, absorbance at OD 600 nm was measured. The results showed that an overnight-incubated *E. coli* culture, when grown for two hours in fresh BHI medium and having an absorbance of 0.1 at OD 600, achieved a concentration of 10^7^ CFU/mL. This suspension was further diluted to achieve a final concentration of ∼100 CFU/mL in sheep blood. The *E. coli* spiked blood cultures showed ∼10^5^ CFU/mL at two hours of incubation and reached 10^6^ CFU/mL after four hours of incubation. Similarly, the *S. aureus* blood culture showed ∼10^2^ CFU/mL at two hours of incubation and reached 10^4^ CFU/mL after four hours of incubation.

### 250 U of SAN showed comparable host depletion to the higher concentration of 500 U

The qPCR performed on the *E. coli* and *S. aureus* DNA extracted after host depletion using two different concentrations of both HL-SAN and M-SAN showed that 250 U (10 μL) of SAN showed similar results compared to 500 U (20 μL) regarding the depletion of host DNA and recovery of bacterial DNA.

For *E. coli* blood cultures (Figure 3A), HL-SAN at both concentrations (250 and 500 U) yielded very similar Ct values (24.5 v/s 24.9) for the host primers, indicating similar host DNA depletion. In comparison, the average bacterial Ct values observed were 24.9 at 250 U and 25.5 at 500 U, indicating a higher recovery of the bacterial DNA at lower SAN concentration (250 U). M-SAN showed similar average Ct values of 23 for the bacterial primers and 22 for the host primers at both the tested concentrations. Compared to HL-SAN, the Ct values for both bacterial and host primers were lower in the samples treated with M-SAN. The *E. coli* samples extracted with the standard BiOstic protocol without any depletion showed lower average Ct values of 21.3 for bacteria and 19.3 for the host (Figure 3A).

**Figure 3.**
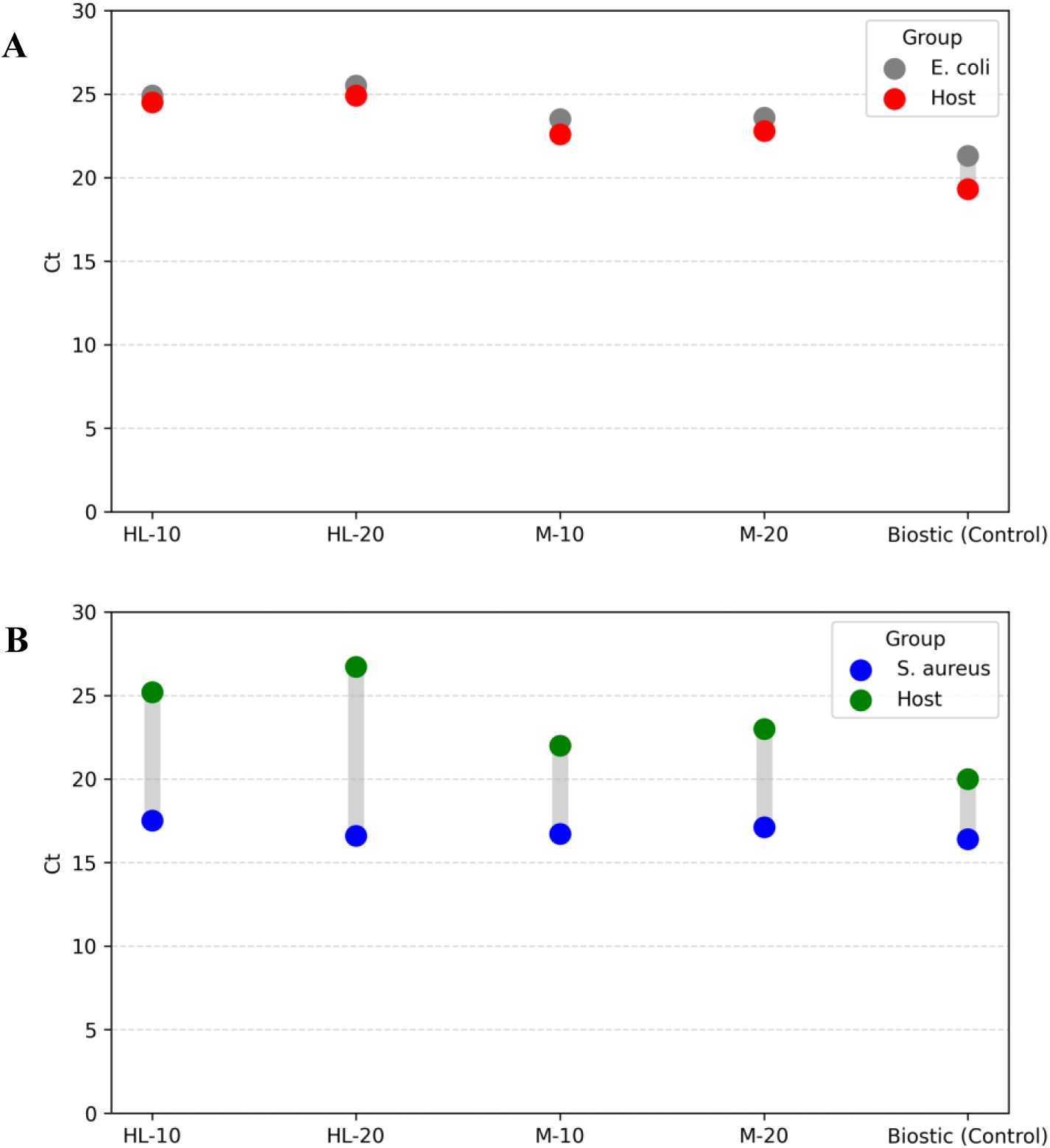
Effect of different enzyme concentrations on host DNA depletion evaluated using qPCR Ct values. (A) *Escherichia coli* (B) *Staphylococcus aureus*. HL-10/20 = 10/20 μL (250/500 U) HL-SAN, M- 10/20 = 10/20 μL (250/500 U) M-SAN In *E. coli* blood cultures, a higher yield of DNA was obtained when a lower concentration of HL- SAN and M-SAN (250 U) was used for host depletion compared to the higher enzyme concentration (500 U). A similar high DNA yield was observed in *S. aureus* samples when M-SAN was used for depletion. However, in *S. aureus* HL-SAN samples, a higher enzyme concentration (500 U) resulted in increased DNA yield (Supplementary Table 2).

In *S. aureus* blood cultures, HL-SAN and M-SAN showed comparable average Ct values of approximately 17 for bacterial primers at both the tested concentrations. However, the Ct values for host primers were 26.7 and 25.2 in HL-SAN at 500 and 250 U, respectively, indicating slightly better host depletion at higher nuclease concentration. Comparable results were obtained for M- SAN, with higher enzyme concentrations yielding elevated host Ct values (average Ct = 23) relative to lower concentrations (average Ct = 22). Overall, both tested enzymes have lower Ct values for bacterial primers compared to host primers, suggesting efficient host depletion. The host depletion was assessed in comparison to the reference BiOstic kit protocol, which involved no depletion. The host Ct values for the non-depleted samples extracted using only the BiOstic kit were lower (average Ct = 20), indicating that more host DNA was present (Figure 3B).

### SAN nuclease activity remained stable across various concentrations of NaCl and MgCl2

**2 and 2.5 M NaCl with 50 mM MgCl2:** *E. coli* blood cultures depleted with HL-SAN and M- SAN, using 2 M NaCl in the buffer, showed less depletion (5–10-fold) compared to those using 2.5 M NaCl in the buffer. In case of HL-SAN at 2 M NaCl, a noticeable loss of the bacterial DNA (2 – 3-fold) was also observed compared to when 2.5 M NaCl was used in the buffer. M-SAN showed similar bacterial DNA recovery at both the tested salt concentrations (Figure 4A). In *S. aureus* samples, HL-SAN provided similar host depletion at both salt concentrations (2 and 2.5 M NaCl). However, M-SAN showed slightly better host depletion (∼5-fold) at 2 M NaCl compared to 2.5 M NaCl (Figure 4B). Both HL-SAN and M-SAN showed higher recovery of bacterial DNA (1–2 fold) at 2 M NaCl compared to 2.5 M NaCl. The DNA yield obtained using M-SAN was comparatively higher than that of HL-SAN (Supplementary Table 3).

**Figure 4.**
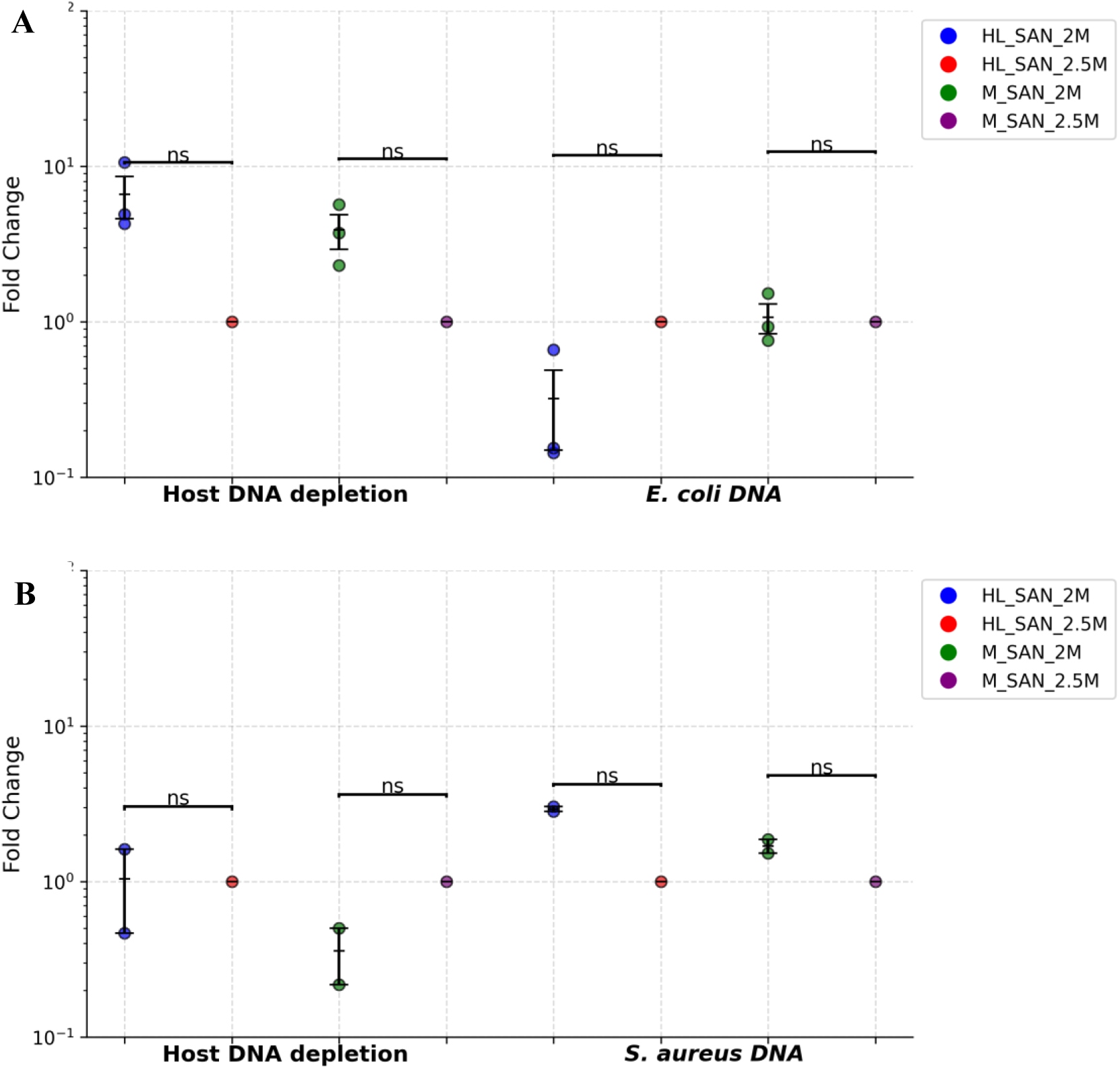
The effect of NaCl concentrations on HL-SAN and M-SAN activity for host DNA depletion. The host DNA depletion and bacterial DNA enrichment are measured as fold change. The 2.5 M NaCl concentration is used as a reference and compared with the 2 M NaCl concentration. The concentration of MgCl2 is maintained at a constant 50 mM (A) *Escherichia coli* and (B) *Staphylococcus aureus*. 2M/2.5M = NaCl

**15 and 50 mM MgCl2 with 2.5 M NaCl:** Different concentrations of MgCl2 (15 and 50 mM) were used as part of the buffer, while maintaining a constant NaCl concentration of 2.5 M (Supplementary Table 4). HL-SAN provided similar host depletion and bacterial DNA recovery at both tested MgCl2 concentrations in *E. coli* blood cultures. The M-SAN-depleted *E. coli* samples showed slightly more host depletion (2–3-fold) at 15 mM MgCl2 compared to 50 mM MgCl2; however, bacterial DNA recovery was also lower (1–2-fold) at 15 mM MgCl2 (Figure 5A). In the case of *S. aureus* blood cultures, both HL-SAN yielded similar host DNA depletion and bacterial DNA recovery at 15 mM and 50 mM MgCl2 (Figure 5B). However, M-SAN at 15 mM MgCl2 showed 1-2 fold higher host DNA depletion compared to 50 mM MgCl2 (Figure 5B). Overall, no significant difference was observed in host DNA depletion or bacterial DNA recovery for both enzymes at different MgCl2 concentrations in *E. coli* and *S. aureus* blood cultures.

**Figure 5.**
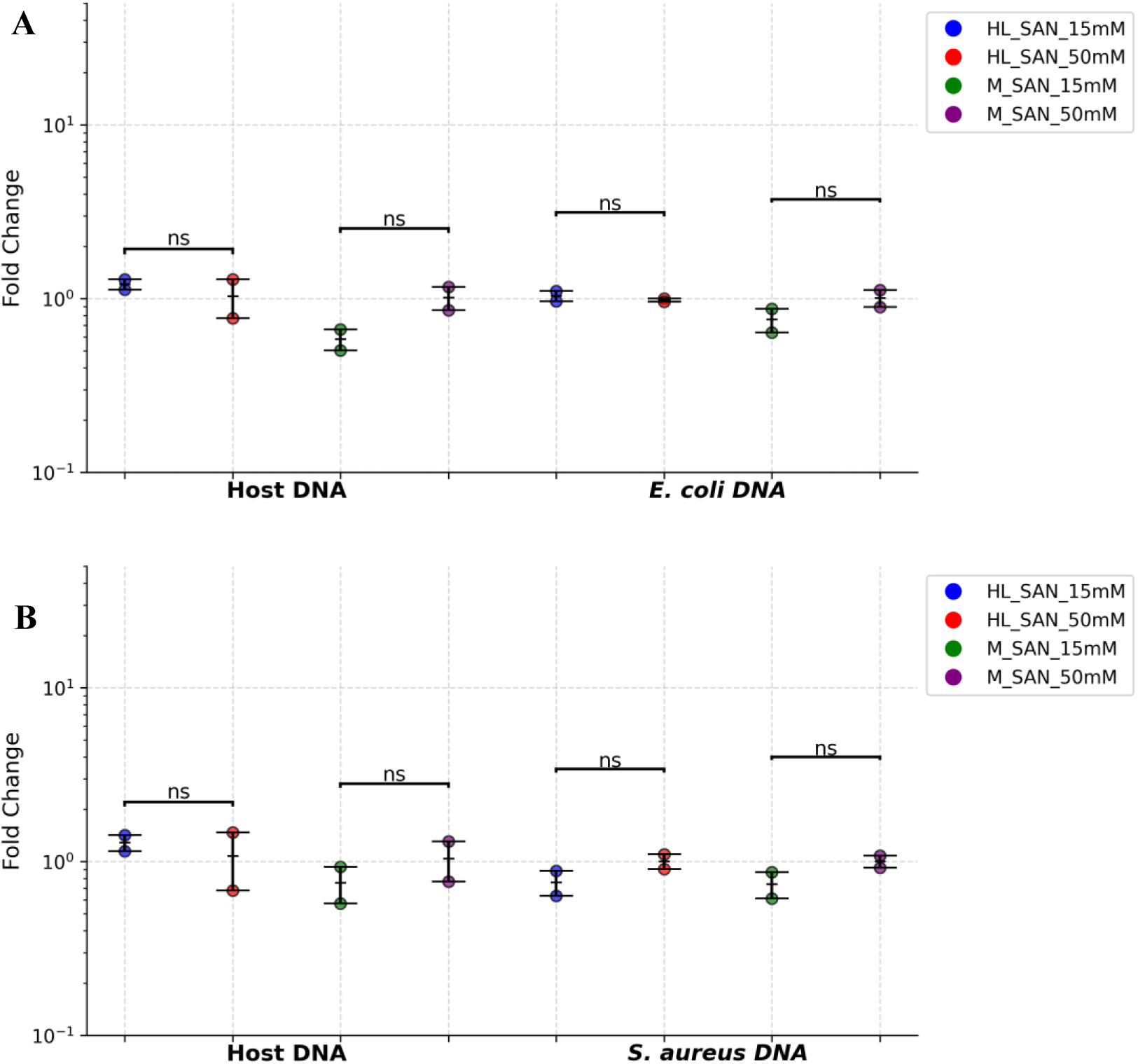
The effect of MgCl2 concentrations on HL-SAN and M-SAN activity for host DNA depletion. The host DNA depletion and bacterial DNA enrichment are measured as fold change. The 50 mM MgCl2 concentration is used as a reference for comparison with the 15 mM MgCl2. The concentration of NaCl is kept constant at 2.5 M (A) *Escherichia coli* and (B) *Staphylococcus aureus*. 15mM/50mM = MgCl2

**0.5 M NaCl and 15 mM MgCl2 vs 2.5 M NaCl and 50 mM MgCl2:** After further reducing the concentration of NaCl to 0.5 M and maintaining the MgCl2 concentration at 15 mM, both enzymes were tested for host depletion on the same *E. coli* and *S. aureus* blood cultures (Supplementary Table 5). The results from these concentrations were compared by using 2.5 M NaCl and 50 mM MgCl2 as a reference control. In HL-SAN *E. coli,* the host DNA depletion was slightly less, and the bacterial DNA recovery was similar at lower salt concentrations compared to the reference. *E. coli* samples with M-SAN showed slightly more host depletion (average ΔCt = 0.4) at lower salt concentrations; however, a minor loss of *E. coli* DNA (average ΔCt = 1.2) was also observed (Figure 6A). *S. aureus* blood cultures depleted with HL-SAN showed similar host depletion, but the bacterial DNA recovery was slightly lower (average ΔCt = 0.7) at 0.5 M NaCl and 15 mM MgCl2. Similarly, *S. aureus* with M-SAN exhibited relatively similar host depletion and bacterial DNA recovery at both salt concentrations tested (Figure 6B).

**Figure 6.**
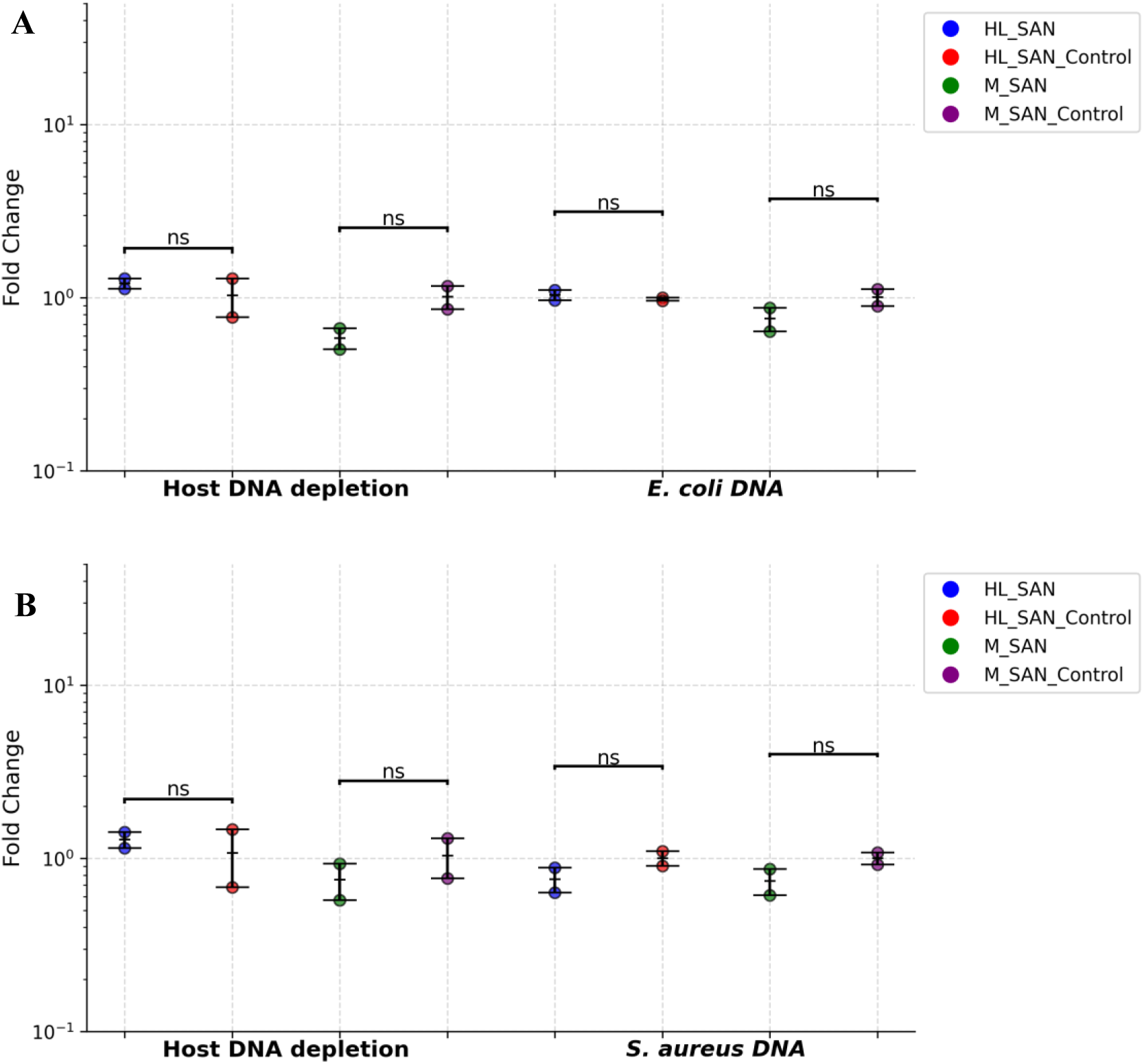
The effect of changing concentrations of NaCl and MgCl2 on HL-SAN and M-SAN activity for host DNA depletion. The host DNA depletion and bacterial DNA enrichment are measured as fold change. The 2.5 M NaCl and 50 mM MgCl2 concentrations are used as a reference control for comparison with 0.5 M NaCl 15 mM MgCl2. (A) *E*. *coli* (B) *S. aureus*. HL_SAN/M_SAN = 0.5 M NaCl and 15 mM MgCl2, HL_SAN/M_SAN control = 2.5 M NaCl and 50 mM MgCl2

### 6 minutes of bead beating improved the recovery of longer DNA fragments

To evaluate the impact of reduced bead-beating duration (6 minutes) on DNA fragment size and nanopore sequencing performance, blood cultures of *E. coli* and *S. aureus* were processed using an identical DNA extraction protocol, except for the duration of the BB. In the modified protocol, the standard 10-minute continuous BB was replaced with three cycles of 2-minute BB, each followed by a 1-minute incubation on ice. The extracted DNA samples were run on the TapeStation and compared. It was observed that reduced bead beating improved the recovery of longer DNA fragments. *E. coli* sample processed with HL-SAN, following the standard 10 minutes of BB, showed the most significant DNA fragmentation with fragments of 2110 bp and concentration of 0.104 ng/μL (Figure 7A). However, when the BB time was reduced, a longer DNA fragment of 11604 bp (0.234 ng/μL) was obtained (Figure 7B). With M-SAN, the *E. coli* samples yielded a relatively larger DNA fragment of 15209 bp at 10 minutes of BB compared to the 13210 bp fragment obtained at reduced BB (6 minutes). However, the concentration of the DNA fragment (10.2 ng/μL) was still higher at 6-minute BB compared to the 10-minute BB (3.23 ng/μL) (Figure 7).

**Figure 7.**
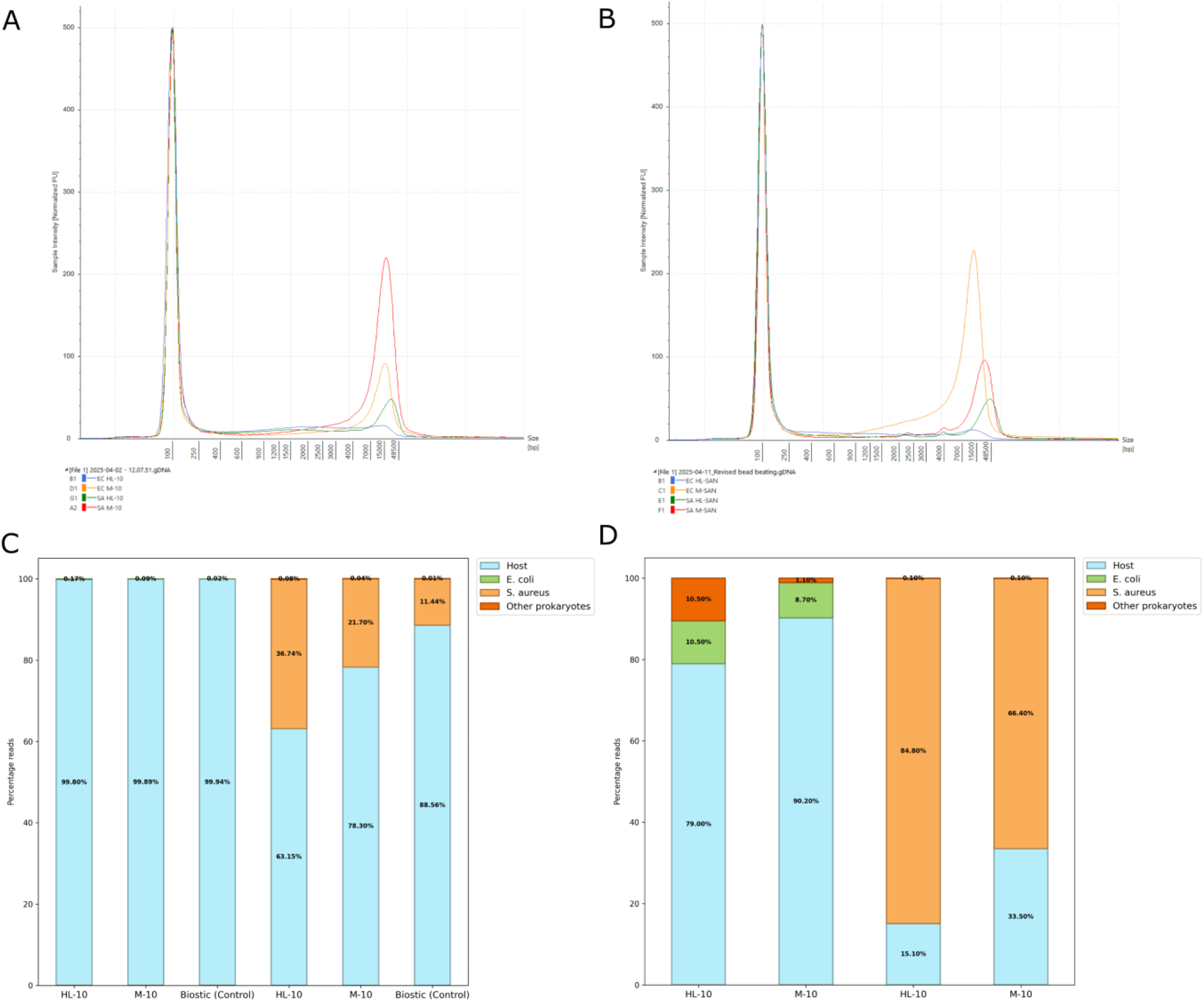
The DNA fragment lengths analyzed using a TapeStation, and the relative distribution of sequencing reads generated by nanopore sequencing from blood cultures. (A) 10 minutes BB (B) 6 minutes BB (C) 10 minutes BB (D) 6 minutes BB. EC = *Escherichia coli*, SA = *Staphylococcus aureus*.

*S. aureus* samples with HL-SAN exhibited two distinct 1717 and 21401 bp DNA fragments at 10 minutes of BB (Figure 7A). However, the samples with 6-minute BB yielded three longer DNA fragments of 2251, 4014, and 41981 bp (Figure 7B). Similarly, *S. aureus* processed with M-SAN also showed longer DNA fragments of 2278, 4010, and 23739 bp with 6-minute BB compared to the DNA fragment (16793 bp) obtained with 10-minute BB (Figure 7).

The sequencing results also showed that, when using the default 10-minute BB time, the number of target pathogen reads recovered was lower compared to using a shorter BB time (6 minutes). In *E. coli* blood cultures using HL-SAN, M-SAN, and the BiOstic kit alone, followed by 10 minutes of BB, most of the extracted DNA was mapped against the host genome. The *S. aureus* samples yielded a higher percentage of bacterial reads. *S. aureus* samples processed with HL-SAN, M- SAN, and BiOstic alone yielded 36%, 22%, and 11% reads, respectively, that mapped to the target pathogen, while the remaining reads aligned with the host genome (Figure 8A). However, when the BB time was reduced to 6 minutes, the number of bacterial reads recovered from nanopore sequencing increased significantly. *E. coli* blood cultures yielded 10.5% (HL-SAN) and 8.7% (M- SAN) of the total reads mapping to the target pathogen when the 6-minute BB protocol was employed. Similarly, in *S. aureus* samples, reducing the bead beating duration resulted in an increased proportion of pathogen-specific reads, with 85% and 66% of total reads mapping to the target organism in HL-SAN and M-SAN treated samples, respectively (Figure 8B).

## Discussion

Rapid diagnosis of BSIs and sepsis is crucial to reducing mortality and the spread of antibiotic resistance. However, the current methods for sepsis diagnosis are based on traditional blood cultures, followed by biochemical tests, MALDI-ToF-MS, or qPCR-based detection [36]. These methods can take up to 5 days for the identification of the causative pathogens and the associated antibiotic resistance [37]. Nanopore sequencing has the potential to overcome these challenges by providing fast and accurate identification of pathogens and antibiotic resistance genes [15,24,34,35,38]. The most crucial step in sequencing-based diagnosis of bloodstream infections is the extraction of microbial DNA and reducing the contamination of host DNA [39]. In this study, we aimed to develop a saponin-based selective lysis method for host cell lysis and utilize endonucleases, such as HL-SAN and M-SAN, to remove the released host DNA, followed by the extraction of bacterial DNA. Additionally, we tested various concentrations of these enzymes and different salt levels to determine the optimal conditions for depleting the host DNA. The results showed that 4% saponin, followed by 10 μL (250 U) of HL-SAN/M-SAN enzymes at a salt concentration of 2.5 M NaCl and 50 mM MgCl2, provided efficient depletion of the host DNA in *E. coli* and *S. aureus* blood cultures.

### 4% saponin is effective for the selective depletion of the host DNA

Chemically, saponins are amphipathic glycosides, characterized by a hydrophobic aglycone (sapogenin) backbone, typically a triterpene or steroid, linked to one or more hydrophilic sugar moieties. Based on this structure, saponins tend to interact more with cholesterol-rich membranes, a feature prevalent in eukaryotic cells rather than bacterial cells [40]. Both planktonic and static growth dynamics in the presence of saponin reveal that saponin more likely disrupts host cells by forming complexes with cholesterol, leading to pore formation in the host cell membrane and allowing osmotic imbalance and leakage of cellular contents. Bacteria that lack cholesterol in their cell walls (which consists mainly of peptidoglycans) are less susceptible to this mechanism, which explains the observed selectivity. This differential membrane composition underpins the targeted action of saponin, making it an optimal agent for host DNA degradation without significant effect on bacterial cells. Previous studies also support this selective lysis of saponin [41,42]. Previous studies have shown that although using saponin, the chromosomal DNA in blood samples was effectively depleted, a large portion of the mitochondrial DNA was still observed [14]. As saponin acts on the cholesterol-rich membranes found in eukaryotic cells, the outer and inner membranes of the mitochondrial cell contain a relatively low number of cholesterol groups [43,44]. Therefore, mitochondrial host cells are usually more resistant to lower concentrations of saponin treatment. *Moragues-Solanas et al*. observed that using 1% saponin for host depletion, around 88% of the total host DNA observed was from mitochondria. However, when the concentration of saponin increased to 3%, the depletion of chromosomal and mitochondrial host DNA was enhanced 10- fold without affecting the bacterial DNA in the samples. These findings are comparable to our results, where we have used 4% saponin and observed that only around 0.1% of the total reads align with the host mitochondrial DNA. The saponin and SAN-based method used in this study effectively depleted the host chromosomal and mitochondrial DNA.

### Lower concentrations of endonucleases provided comparative host depletion to the higher concentrations

In this study, two different concentrations of HL-SAN and M-SAN (250 U and 500 U) are used and compared to see the efficiency of host DNA depletion in blood cultures. The use of optimal enzyme concentration is essential to achieve effective host DNA depletion, as too little enzyme may not be effective, and higher concentrations might adversely affect the downstream extraction process. Previous studies have also employed different enzyme concentrations for testing host DNA depletion in various samples. *Charalampous et al*. initially used 5 + 2 μL of HL-SAN enzyme, and later they optimized host DNA depletion in respiratory samples by using 10 μL of HL-SAN enzyme [21,45]. A similar concentration (10 μL) of HL-SAN was used for the depletion of host DNA in stool, respiratory, and serum samples [45–47]. These studies have demonstrated the effectiveness of host DNA depletion using 10 μL HL-SAN. However, the type of sample used is different, and little work has been done on optimizing the concentration of these enzymes for host DNA depletion in blood samples or cultures.

*Moragues-Solanas et al*. have recently demonstrated that using 10 μL of HL-SAN in combination with saponin-based host cell lysis provides effective host DNA depletion in blood samples spiked with clinically relevant sepsis pathogens, including *E. coli*, *S. aureus*, *K. pneumoniae*, and *E. faecalis* [23]. However, they have not tested different concentrations of the enzymes to see how they will impact the host DNA depletion and later bacterial DNA extraction. In our previous study, we tested the host depletion of HL-SAN and M-SAN using a higher concentration of 20 μL (500 U) on clinical urine samples [24]. The results demonstrated effective depletion of host DNA, which is sufficient for accurately identifying bacterial pathogens and antibiotic resistance genes in clinical urinary tract infection samples using nanopore sequencing. In this study, we have demonstrated that using both enzyme concentrations (250 U and 500 U) provided effective host DNA depletion in *E. coli* and *S. aureus* blood cultures. For *S. aureus,* slightly better host DNA depletion was observed using 500 U (20 μL) of the enzyme compared to 250 U (10 μL). However, in *E. coli* samples, both enzymes provided similar host DNA depletion at both tested concentrations.

Therefore, it can be concluded that although changing the enzyme concentrations has a slight effect on host DNA depletion, the overall host depletion at lower concentrations is still sufficient for downstream processes. The lower concentration of enzyme would reduce the cost per sample by half, as only 10 μL of the SAN enzyme would be used instead of 20 μL. Although the lower enzyme concentration is effective, it is essential to note that this study used spiked blood samples, which may not accurately represent the blood samples from clinical sepsis patients. Typically, in cases of infections such as bloodstream infections or sepsis, the host’s immune response is heightened, resulting in the release of large numbers of leukocytes into the bloodstream [48]. So, the overall host DNA released from leukocytes in sepsis samples would be higher compared to the spiked healthy samples, where no immune response is initiated. Therefore, it is recommended to use higher enzyme concentrations, 500 U (20 μL) of HL-SAN and M-SAN for host DNA depletion in clinical sepsis samples.

### Changing salt concentrations for enzyme activity does not have a significant effect on the host DNA depletion

In this study, we used various salt concentrations as buffers for HL-SAN and M-SAN to investigate whether they would affect enzyme activity and overall host DNA depletion in blood cultures. The results showed that the buffer with higher salt concentrations (2.5 M NaCl and 50 mM MgCl2) provided comparatively better host DNA depletion than buffers with lower salt concentrations. However, the differences in host DNA depletion were not significant when changing the concentrations of NaCl and MgCl2 in the buffers. HL-SAN and M-SAN are salt-active nucleases and can tolerate high levels of salts. The manufacturer (ArcticZymes) suggests a concentration range of 125 mM – 250 mM NaCl and 4 – 15 mM MgCl2 for M-SAN and 400 – 700 mM NaCl and 5 – 50 mM MgCl2 for optimal activity of HL-SAN [49,50]. However, the optimal activity across different sample types, such as blood, urine, and respiratory samples, can vary due to the presence of salt or the absorption of the salt in the buffer by the sample matrix.

Endonucleases catalyze the cleavage of the phosphodiester bonds within the DNA double helix. They introduce single- or double-stranded breaks through hydrolysis of the phosphodiester linkage, explicitly targeting the P–O3′ bond, and generate DNA fragments with 5′-phosphate and 3′-hydroxyl termini [51]. Along with the HL-SAN and M-SAN that we tested in this study, numerous nucleases such as Benzonase (Sigma Aldrich Merck), DNase I, Denarase (c-Lecta), and DNaseMe (Qiagen) are commercially available [52,53]. Most of the nucleases require MgCl2 or CaCl2 as a cofactor for their activity. Nucleases, such as DNase I, are extremely sensitive to high salt concentrations and are unable to perform their activity. Benzonase is one of the most used nucleases in biopharmaceutical production due to its broad substrate specificity and efficiency [54]. Although it exhibits greater tolerance to salt compared to DNase I, elevated salt concentrations can still inhibit its enzymatic activity by interfering with DNA binding or altering the enzyme’s conformation. A salt concentration of 200 mM has been shown to decrease the enzyme activity to less than 10% [55].

In a previous study, different salt concentrations have been used to compare the activity of Benzonase, Denarase, Saltonase, MNase, DnaseMe, M-SAN, and SAN (HL-SAN). These nucleases were tested using salt conditions of 2 mM MgCl2 and higher concentrations recommended by the respective manufacturers. The results indicated that Benzonase, Denarase, MNase, and M-SAN exhibited efficient DNA digestion at MgCl₂ or CaCl₂ concentrations of 2 mM or higher, with MNase requiring CaCl₂ as a cofactor [53]. Benzonase and Denarase have been shown to perform optimally at lower concentrations of NaCl, while M-SAN performed exceptionally well even with higher concentrations of salt. These findings are consistent with our observations in this study, indicating that the use of 15 mM or 50 mM MgCl₂ and varying NaCl concentrations had no significant impact on the enzymatic activity of HL-SAN and M-SAN. Additionally, in our previous study, we demonstrated that HL-SAN and M-SAN effectively depleted host DNA in clinical urine samples using higher salt concentrations of 2.5 M NaCl and 50 mM MgCl2 [24]. In the blood samples of sepsis patients, the background human DNA will be significantly higher due to the leukocytes released into the blood as part of the immune response [56]. Therefore, the optimal activity of nucleases will be beneficial for digesting the released human DNA and subsequently extracting the microbial DNA for downstream diagnostic procedures.

### Reduced bead beating time enhanced the recovery of larger DNA fragments

Nanopore sequencing requires longer DNA fragments for high throughput. Therefore, the integrity of the extracted DNA is crucial for the successful application of nanopore sequencing. After the depletion of host DNA in blood samples through saponin and nucleases, the microbial DNA is extracted. Along with other methods, mechanical lysis using various types of beads is typically employed to lyse microbial cells and release their DNA [57,58]. BB is critical for gram-positive bacteria due to their thick cell walls, which are difficult to lyse using chemical methods [59]. Although BB can enhance nucleic acid yield, excessive BB can lead to DNA shearing, which compromises downstream processes.

In this study, we used the BiOstic kit, which employs BB to lyse the cells. The default kit protocol suggests using 10 minutes of continuous BB. However, we observed that using the 10-minute BB method promoted DNA shearing, which compromised sequencing throughput, especially for gram-negative bacteria such as *E. coli*. The BiOstic kit is designed for DNA extraction directly from blood cultures and does not include any host DNA depletion mechanisms. In this study, the blood culture samples used for extraction were host DNA-depleted using nucleases and thus had little host DNA available. Therefore, the presence of a higher beads-to-cells ratio might have caused fragmentation of the microbial DNA due to extended bead beating.

The bead beating time was then reduced to three cycles, each consisting of two minutes of BB followed by one minute of incubation on ice. Reducing bead beating resulted in improved fragment lengths and increased downstream sequencing throughput compared to the default bead beating. Previously, BB times of 0, 3, 10, 15, and 20 minutes have been investigated on bacterial and fungal communities, and the results showed that 20 minutes of BB is the most appropriate for recovering most of the microbial DNA [58]. These results contrast with those observed in this study and our previous study, where 6 minutes of BB was sufficient for the recovery of high-integrity bacterial DNA for nanopore sequencing [24]. The recovery of high-integrity DNA using BB is also strictly dependent on the type of instrument, the speed, and the type of beads used. Similar to previous studies, we also observed that increasing the bead beating time reduced the recovery of high-integrity DNA from the gram-negative bacterium *E. coli* [60]. Therefore, we suggest that a BB time of 6 minutes will be enough to recover DNA from both gram-positive and gram-negative bacteria.

In summary, these results demonstrated that 4% saponin, followed by digestion with SAN nucleases at a concentration of 250 units (10 μL), was an efficient method for selectively depleting host DNA in blood cultures. Additionally, the varying concentrations of NaCl and MgCl2 as a buffer for enzymes had no significant effect on enzyme performance or host DNA depletion. The reduced BB time of 6 minutes is recommended as it provides highly intact DNA and longer fragments compared to the default 10-minute BB from the BiOstic kit. Furthermore, the optimized method in this study is compatible with downstream processes such as qPCR and nanopore sequencing. The results from this study provided helpful information for the depletion of host DNA using saponin and SAN enzymes. However, in this study, the host DNA depletion was assessed using simulated (bacterial-spiked) blood cultures. These samples may not accurately represent actual sepsis patient samples. Also, the number of bacteria tested in this study was limited to only *E. coli* and *S. aureus*. Therefore, in future studies, more sepsis-relevant pathogens must be tested, along with testing on the actual clinical samples. We are currently testing the protocol using actual clinical sepsis samples.

## Acknowledgements

This study was supported by the OH-AMR-Diag project, funded by the Research Council of Norway (project number 336420).

## Author Contributions

**Resources:** Rafi Ahmad. **Conceptualization:** Rafi Ahmad and Jawad Ali. **Methodology:** Jawad Ali, Anurag Basavaraj Bellankimath, Jonathan Hira, and Crystal Chapagain. **Investigation and formal analysis:** Jawad Ali, Anurag Basavaraj Bellankimath, Jonathan Hira, and Rafi Ahmad. **Supervision:** Rafi Ahmad. **Writing – original draft:** Jawad Ali and Jonathan Hira. **Writing – review & editing:** Jawad Ali, Jonathan Hira, Anurag Basavaraj Bellankimath, and Rafi Ahmad.

