## Supplementary Figure 1 for "Development and optimization of the host DNA depletion in blood cultures using a saponin and SAN nucleases-based method"

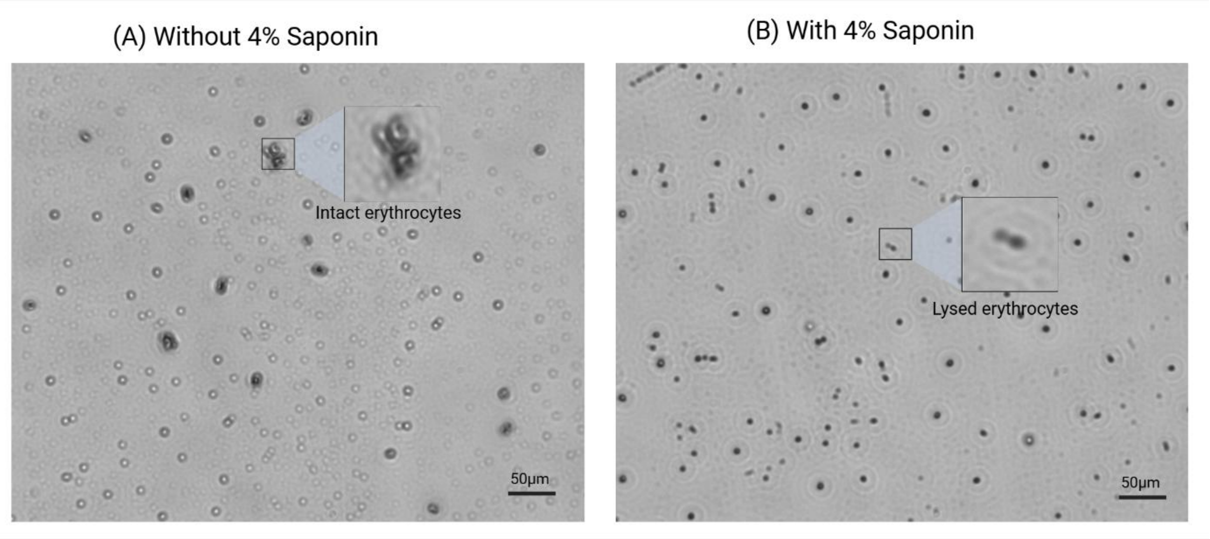


**Fig S1:** Illustrates the targeted cellular lytic effect of saponin at 4% concentration. (A) Without saponin, healthy intact erythrocytes are observed, but lysed erythrocytes are prominent in the presence of saponin.
