## Supplementary Table 1 for "Development and optimization of the host DNA depletion in blood cultures using a saponin and SAN nucleases-based method"

**Supplementary Table 1:** Information about the primers used for the qPCR amplification of host (sheep), *E. coli* and *S. aureus* DNA from blood cultures.

| **Target species** | **Target gene** | **Amplicon size (bp)** | **Primer sequences** | **References** |
|---|---|---|---|---|
| Sheep | *HPRT1* | 102 | FP: CACTGGGAAGACAATGCAGA<br>RP: ACACTTCGAGGGGTCCTTTT | [1] |
| *E. coli* | *UspA* | 850 | FP: CCGATACGCTGCCAATCAGT<br>RP: ACGCAGACCGTAGGCCAGAT | [2] |
| *S. aureus* | *Nuc* | 65 | FP: GGGTTGATACGCCAGAAACG<br>RP: TGATGCTTCTTTGCCAAATGG | [3] |
