## Supplementary Table 2 for "Development and optimization of the host DNA depletion in blood cultures using a saponin and SAN nucleases-based method"

**Supplementary Table 2:** Host DNA depletion using different concentrations of HL-SAN and M-SAN, followed by extraction with the BiOstic kit. HL-10/20 = 10/20 µL (250/500 U) HL-SAN, M-10/20 = 10/20 µL (250/500 U) M-SAN

| **Samples** | **Endonuclease used** | **Enzyme units (U)** | **DNA yield (ng)** | **Nanodrop quality check** | | **Average size (bp) Tapestation** | **DIN value** | **Average CT (Bacteria)** | **Average CT (Host)** |
|---|---|---|---|---|---|---|---|---|---|
| | | | | **260/280** | **260/230** | | | | |
| *E. coli* | HL-SAN | 250 | 55 | 1.4 | 0.3 | 4007 | 3.8 | 24.9 | 24.5 |
| | HL-SAN | 500 | 45 | 1.4 | 0.3 | 2809 | 3.1 | 25.5 | 24.9 |
| | M-SAN | 250 | 105 | 1.5 | 0.2 | 9792 | 7 | 23.5 | 22.6 |
| | M-SAN | 500 | 90 | 1.5 | 0.5 | 6874 | 6.6 | 23.6 | 22.8 |
| | No depletion (Biostic kit only) | - | 1000 | 1.7 | 0.9 | 7321 | 6.4 | 21.3 | 19.3 |
| *S. aureus* | HL-SAN | 250 | 75 | 1.3 | 0.1 | 8638 | 6.2 | 17.5 | 25.2 |
| | HL-SAN | 500 | 115 | 1.3 | 0.5 | 13632 | 6.6 | 16.6 | 26.7 |
| | M-SAN | 250 | 230 | 1.6 | 1.1 | 12536 | 7.3 | 16.7 | 22 |
| | M-SAN | 500 | 145 | 1.6 | 0.3 | 14575 | 7.5 | 17.1 | 23 |
| | No depletion (Biostic kit only) | - | 800 | 1.7 | 1.7 | N/A | N/A | 16.4 | 20 |
