## Supplementary Table 3 for "Development and optimization of the host DNA depletion in blood cultures using a saponin and SAN nucleases-based method"

**Supplementary Table 3:** Host depletion and DNA extraction from blood cultures using 2 and 2.5 M NaCl and 50 mM $MgCl_2$ (constant) as buffer for enzyme activity.

| Samples | Endonuclease used | Replicates | NaCl Conc (M) | DNA yield (ng) | Nanodrop quality check | | Average CT (Bacteria) | Average CT (Host) |
|---|---|---|---|---|---|---|---|---|
| | | | | | 260/280 | 260/230 | | |
| *E. coli* | HL-SAN | 1 | 2 | 378 | 1.52 | 0.94 | 25.5 | 21.1 |
| | | 2 | 2 | 218 | 1.24 | 0.83 | 27.7 | 22.2 |
| | | 3 | 2 | 235 | 1.18 | 0.78 | 27.6 | 22.4 |
| | | | 2.5 | 55 | 1.4 | 0.3 | 24.9 | 24.5 |
| | M-SAN | 1 | 2 | 520 | 1.44 | 0.76 | 22.9 | 20.1 |
| | | 2 | 2 | 300 | 1.36 | 0.86 | 23.6 | 20.7 |
| | | 3 | 2 | 190 | 1.29 | 0.63 | 23.9 | 21.4 |
| | | | 2.5 | 105 | 1.5 | 0.2 | 23.5 | 22.6 |
| *S. aureus* | HL-SAN | 1 | 2 | 210 | 1.44 | 1.04 | 15.9 | 26.3 |
| | | 2 | 2 | 235 | 1.54 | 1 | 16 | 24.5 |
| | | | 2.5 | 75 | 1.3 | 0.1 | 17.5 | 25.2 |
| | M-SAN | 1 | 2 | 216 | 1.42 | 0.8 | 16.1 | 24.2 |
| | | 2 | 2 | 291 | 1.51 | 0.81 | 15.8 | 23 |
| | | | 2.5 | 230 | 1.6 | 1.1 | 16.4 | 20 |
