## Supplementary Table 4 for "Development and optimization of the host DNA depletion in blood cultures using a saponin and SAN nucleases-based method"

**Supplementary Table 4:** Host depletion and DNA extraction from blood cultures using 2.5 M NaCl (constant) and 15 and 50 mM $MgCl_2$ as buffer for enzyme activity.

| **Samples** | **Endonuclease used** | **Replicates** | **$MgCl_2$ conc (mM)** | **DNA yield (ng)** | **Average CT (Bacteria)** | **Average CT (Host)** |
|---|---|---|---|---|---|---|
| *E. coli* | HL-SAN | 1 | 15 | 172 | 19.3 | 22.9 |
| | | 2 | 15 | 145 | 19.6 | 23.1 |
| | | 1 | 50 | 157 | 19.5 | 22.9 |
| | | 2 | 50 | 111 | 19.6 | 23.6 |
| | M-SAN | 1 | 15 | 176 | 19.1 | 22.1 |
| | | 2 | 15 | 207 | 18.6 | 21.7 |
| | | 1 | 50 | 330 | 18.3 | 20.8 |
| | | 2 | 50 | 286 | 18.6 | 21.3 |
| *S. aureus* | HL-SAN | 1 | 15 | 198 | 19.7 | 22.5 |
| | | 2 | 15 | 235 | 20.2 | 22.2 |
| | | 1 | 50 | 307 | 19.4 | 22.2 |
| | | 2 | 50 | 175 | 19.7 | 23.3 |
| | M-SAN | 1 | 15 | 268 | 20.2 | 21.3 |
| | | 2 | 15 | 177 | 20.7 | 22 |
| | | 1 | 50 | 358 | 19.8 | 20.8 |
| | | 2 | 50 | 168 | 20.1 | 21.6 |
