## Supplementary Table 5 for "Development and optimization of the host DNA depletion in blood cultures using a saponin and SAN nucleases-based method"

**Supplementary Table 5:** Host depletion and DNA extraction from blood cultures using 0.5 M NaCl and 15 mM $MgCl_2$ as buffer for enzyme activity.

| Samples | Host depletion | NaCl conc (M) | $MgCl_2$ conc (mM) | Replicates | DNA yield (ng) | Average CT (Bacteria) | Average CT (Host) |
|---|---|---|---|---|---|---|---|
| *E. coli* | HL-SAN | 0.5 | 15 | 1 | 194 | 19.3 | 22.4 |
| | | | | 2 | 255 | 19.3 | 21.6 |
| | | 2.5 | 50 | 1 | 157 | 19.5 | 22.9 |
| | | | | 2 | 111 | 19.6 | 23.6 |
| | M-SAN | 0.5 | 15 | 1 | 309 | 19.5 | 21.5 |
| | | | | 2 | 295 | 19.7 | 21.4 |
| | | 2.5 | 50 | 1 | 330 | 18.3 | 20.8 |
| | | | | 2 | 286 | 18.6 | 21.3 |
| *S. aureus* | HL-SAN | 0.5 | 15 | 1 | 207 | 20.2 | 22.3 |
| | | | | 2 | 183 | 20.1 | 22.5 |
| | | 2.5 | 50 | 1 | 307 | 19.4 | 22.2 |
| | | | | 2 | 175 | 19.7 | 23.3 |
| | M-SAN | 0.5 | 15 | 1 | 395 | 19.7 | 20.6 |
| | | | | 2 | 486 | 19.8 | 20.7 |
| | | 2.5 | 50 | 1 | 358 | 19.8 | 20.8 |
| | | | | 2 | 168 | 20.1 | 21.6 |
